# A dopaminergic basis of behavioral control

**DOI:** 10.1101/2024.09.17.613524

**Authors:** Ian C. Ballard, Daniella J. Furman, Anne S. Berry, Robert L. White, William J. Jagust, Andrew S. Kayser, Mark D’Esposito

## Abstract

Both goal-directed and automatic processes shape human behavior, but these processes often conflict. *Behavioral control* is the decision about which process guides behavior. Despite the importance of behavioral control for adaptive decision-making, its neural mechanisms remain unclear. Critically, it is unknown if there are mechanisms for behavioral control that are distinct from those supporting the formation of goal-relevant knowledge. We performed deep phenotyping of individual dopamine system function by combining multiple PET scans, fMRI, and dopaminergic drug administration in a within-subject, double-blind, placebo-controlled design. Subjects performed a rule-based response time task, with goal-directed and automatic decision-making operationalized as model-based and model-free influences on behavior. We found a double dissociation between two aspects of ventral striatal dopamine physiology: D2/3 receptor availability and dopamine synthesis capacity. Convergent and causal evidence indicated that D2/3 receptors regulate behavioral control by enhancing model-based and blunting model-free influences on behavior but do not affect model-based knowledge formation. In contrast, dopamine synthesis capacity was linked to the formation of model-based knowledge but not behavioral control. D2/3 receptors also modulated frontostriatal functional connectivity, suggesting they regulate behavioral control by gating prefrontal inputs to the striatum. These results identify central mechanisms underlying individual and state differences in behavioral control and point to striatal D2/3 receptors as targets for interventions for improving goal-directed behavior.

## Introduction

Behavior is controlled by deliberate and automatic processes^1^, often conceptualized as “model-free” and “model- (or rule-) based decision making, respectively^2^. These two systems can conflict, such as when the desire to purchase a pint of ice cream in the supermarket conflicts with one’s goal to consume less sugar. The process of determining which system governs behavior, or deciding how to decide, is termed *behavioral control*^*3*,*4*^. Inappropriate behavioral control has a significant societal impact, from responding to phone notifications while driving to making poor retirement savings decisions^5^. However, the neural mechanisms underlying behavioral control remain unclear.

Emerging evidence points to a role for striatal dopamine D2 receptors in behavioral control^6^. Theoretical models of behavioral control suggest that negative outcomes should cause decision-makers to rely more strongly on model-based information^4,7,8^. Striatal D2-expressing neurons appear well-suited to perform this function. D2-expressing neurons causally update future behavior in response to negative outcomes^9–11^. Moreover, D2-receptors modulate the influence of prefrontal inputs onto striatal neurons^12–15^. Therefore, striatal D2-receptor activation may facilitate the use of model-based knowledge. However, evidence that increased D2 receptor availability or D2 receptor activation enhances model-based control is lacking.

Behavioral control depends on sufficient learning of model-based and model-free knowledge: one cannot decide to use model-based knowledge that does not exist. Distinct aspects of striatal dopamine physiology may influence model-free learning, model-based learning, and behavioral control. There is strong evidence that reward prediction errors conveyed by a subset of dopamine neurons underlie model-free learning^16–18^. Additionally, presynaptic dopamine synthesis capacity in the striatum is linked to cognitive processes important for model-based learning, including the willingness to engage in cognitive effort^19,20^ and working memory capacity^21^. We sought to distinguish the roles of dopamine synthesis capacity and D2/3 receptor function in behavioral control versus the acquisition of model-based knowledge.

In the same human subjects, we employed pharmacological dopamine manipulations, PET imaging of multiple aspects of dopamine system physiology, functional fMRI probes of frontostriatal circuits, and a behavioral task that distinguishes rule-based from model-free contributions to behavior. We found a double dissociation between the striatal D2/3 function and dopamine synthesis capacity in their influence on behavioral control and the formation of rule knowledge, respectively. Additionally, neuroimaging analyses revealed that striatal D2/3 receptors modulated frontostriatal functional connectivity in a qualitatively similar manner as behavioral control. These results identify a specific role for striatal D2/3 function in regulating behavioral control in humans.

## Results

Healthy subjects (n = 77, 48 female sex, median age = 21, age range 18 - 30, standard deviation = 2.46 years) performed a speeded reaction-time task to earn rewards following ingestion of a placebo, bromocriptine (a D2 agonist), and tolcapone (a catechol-O-methyltransferase [COMT] inhibitor) in a within-subject, double-blind study (Figure 1A). In the task, targets were preceded by reward-predictive stimuli. One of the stimulus features was associated with a higher probability of reward (Figure 1B). Additionally, faster responses on rewarded trials led to larger reward outcomes. Therefore, subjects could prepare faster responses for reward predictive stimuli to earn higher rewards. Subjects responded more quickly to high-reward than low-reward stimuli across drug conditions, *Z =* -7.8, *p* < .001, *β* = -2.0 ms (Figure 1C). Additionally, subjects developed explicit knowledge of the task rules: subjects rated the high-reward stimuli as having a higher reward probability than low-reward stimuli, *Z =* 52, *p* < .001, *β* = 21% (Figure 1D).

**Figure 1.**
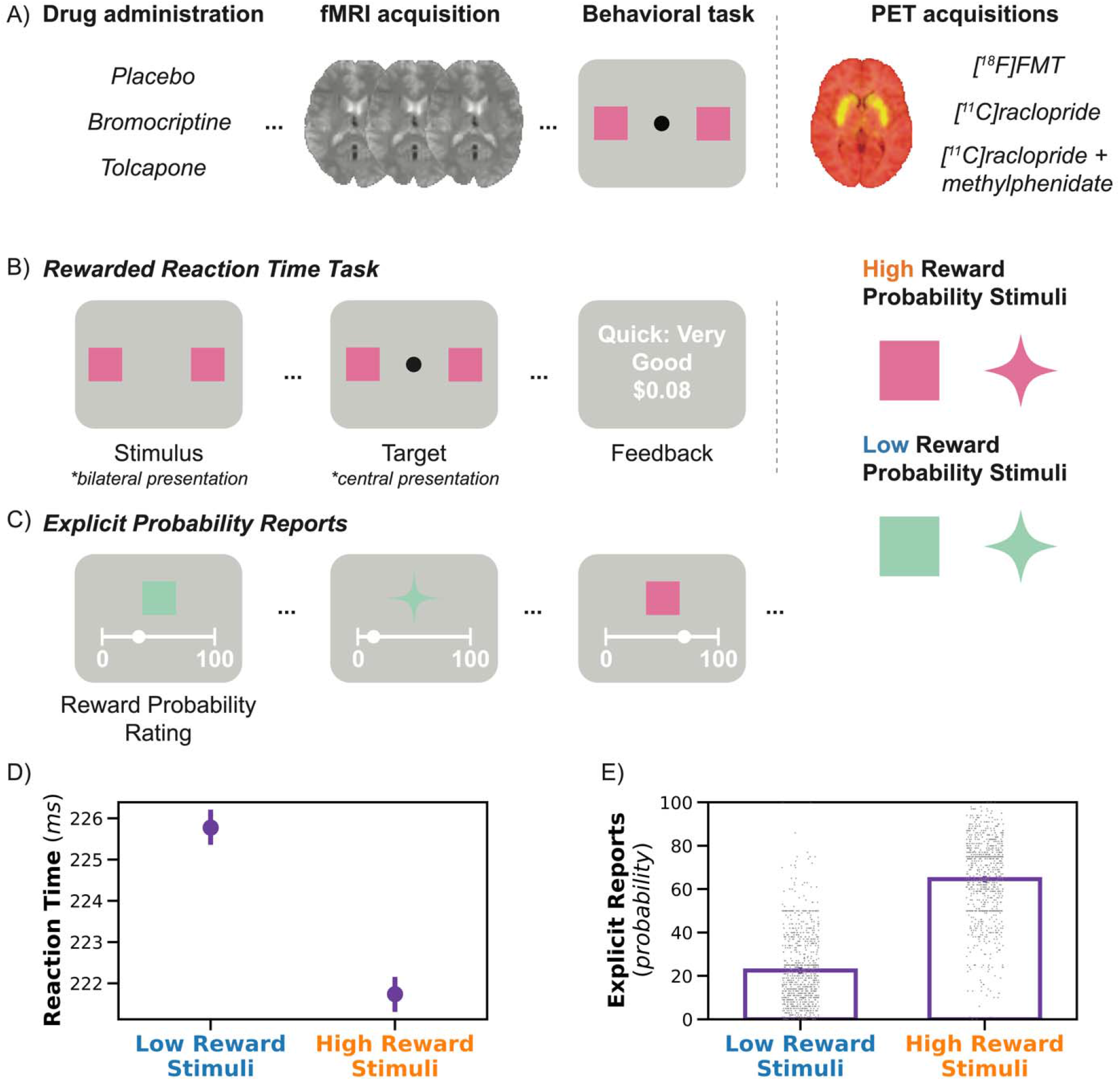
Experimental design and behavioral performance. **(A)** In a double-blind and placebo-controlled study, subjects (*n* = 77) came to the lab for three drug sessions (placebo, bromocriptine, and tolcapone). Subjects then underwent fMRI imaging before completing the behavioral task outside the scanner. On different days, a subset of the subjects underwent PET neuroimaging. **(B)** Subjects performed a speeded reaction time task with probabilistic rewards. Each trial began with a stimulus presented on the left and right side of the screen. The stimulus features varied in color and shape, and one of the features indicated high reward probability (“pink” in the example). After a variable interval, a target appeared centrally. Subjects responded to the target by pressing a key as quickly as possible. On rewarded trials, subjects received larger rewards for faster responses. **(C)** Subjects reported the reward probability of each of the four stimuli by moving a slider to indicate the probability of reward associated with each stimulus. They made these judgments once halfway through the task and once at the end of the task. **(D)** Reaction times were faster for high-reward probability stimuli than for low-reward probability stimuli. **(E)** Explicit probability reports were larger for high-than low-reward probability stimuli. *Error bars depict the standard error of the mean (SEM)*.

Our task design aimed to distinguish the contributions of rule-based and model-free influences on behavior. Rules are abstract representations of task variables (i.e., the relationship between stimulus features and reward probability) that constitute a model of the task. Subjects could calibrate their responses to the stimuli according to the model-free reward value, learned via reinforcement learning^22,23^ (Figure 2A), or according to rule-based knowledge of which feature was reward predictive. Behavioral control refers to the relative contribution of reinforcement learning versus rule-based knowledge to behavior. Importantly, model-free learning is not the most effective response strategy for this task because the rules are fixed. For example, model-free learning reduces the value of a high-reward stimulus after a sporadic reward omission, which would erroneously lead to slower responses upon the next presentation of that stimulus. Humans can learn simple feature-response rules in only a few trials^24^ and could choose to uniformly respond more quickly to high-reward stimuli thereafter (Figure 2B).

**Figure 2.**
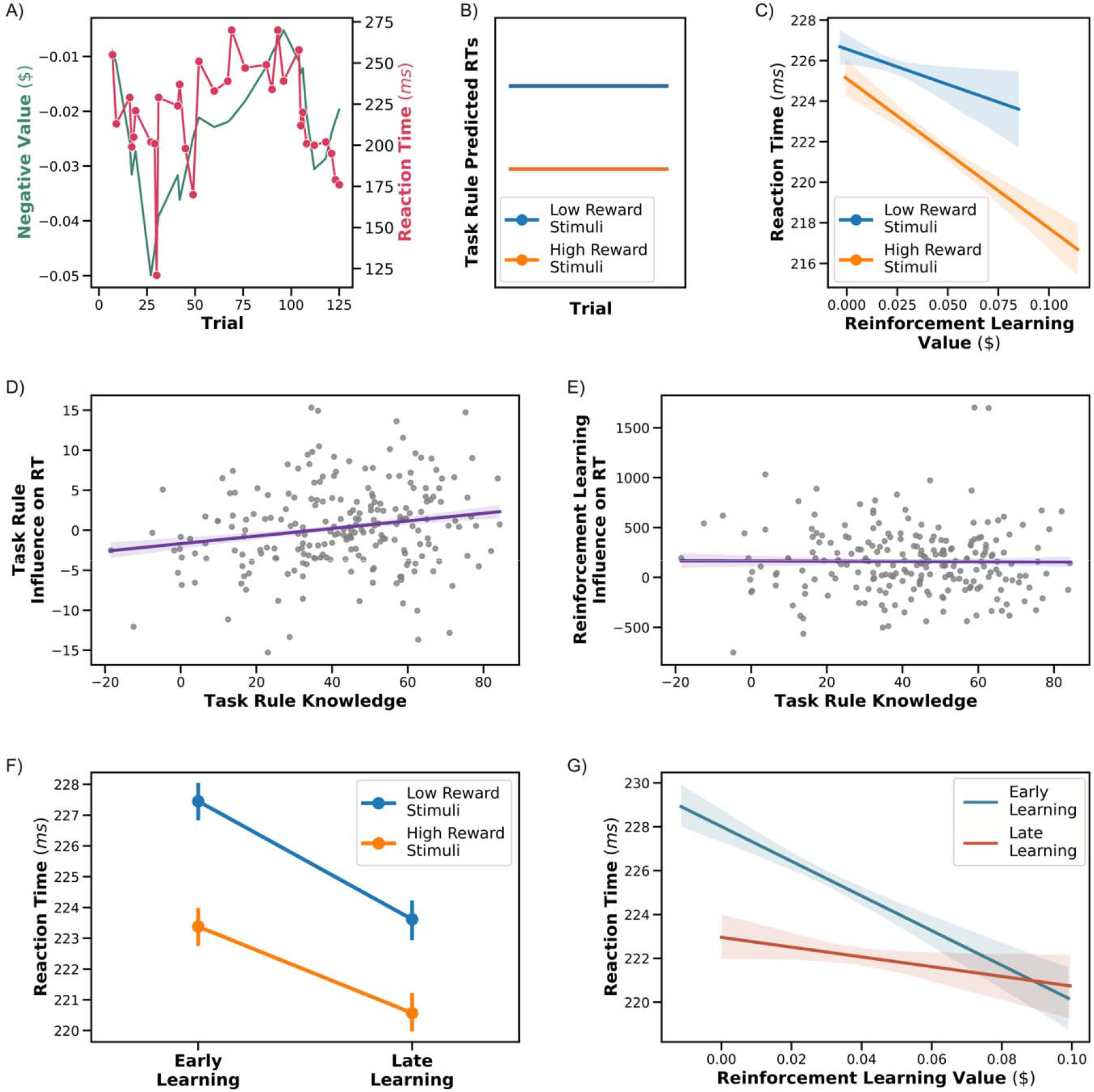
Dissociating the influence of reinforcement learning from task rules on reaction times. **(A)** Example data from one subject showing the trial-by-trial relationship between reinforcement learning value and reaction times. Note that the reinforcement learning values have been inverted for ease of visualization (higher values are associated with *faster* reaction times). **(B)** Predicted reaction times if subjects respond according to task rules. In contrast to reinforcement learning values, task rule values are constant across trials. **(C)** Reaction times are influenced by reinforcement learning, with higher value trials associated with faster reaction times and task rules, with an overall faster reaction time for high reward stimuli. Reinforcement learning values were generated using a cross-validated model-fitting procedure. **(D)** Subjective reports of task rule knowledge are positively related to the influence of task rules on reaction times. The x-axis denotes the difference in the reward probability ratings between high- and low-reward probability stimuli. The y-axis denotes the change in RT in ms between high- and low-reward stimuli. **(E)** In contrast to *C*, subjective reports of task rule knowledge are unrelated to the influence of reinforcement learning on reaction times. **(F)** Task rules influence reaction times in both early and late learning. **(G)** Reinforcement learning values influence reaction times more strongly early in learning. Figure depicts a partial regression that removes task rule differences from reinforcement learning values. *In D and E, dots depict one session from one subject. Error bars depict the SEM*.

We found that reinforcement learning and task rules (i.e., categorical rule effects) were independently associated with reaction times. In a cross-validated analysis (see *Procedures*), reinforcement learning values were associated with faster reaction times, *Z* = -4.0, *p* < .001, *β* = -0.71 ms per penny (Figure 2C). Critically, we also found an independent influence of task rules on reaction times, *Z* = -3.2, *p* = .001, *β* = -1.1 ms (Figure 2C), indicating that subjects used both rule-based and reinforcement-learning-based strategies in the task. Our task is, therefore, well-suited to assess how dopamine physiology influences behavioral control.

We performed several validation checks to determine whether we could reliably distinguish the influence of task rules from reinforcement learning. First, our model performed well in a parameter recovery study (Table S1)^25^. Second, we reasoned that individual variability in task rule knowledge ought to relate to variability in the influence of task rules on reaction times. We operationalized task rule knowledge as the difference in the reward probability ratings between high- and low-reward probability stimuli. Task rule knowledge was correlated with the influence of task rules on reaction times, *Z* = 2.1, *p* = .038, *β* = 1.0 (Figure 2D). In contrast, there was no relationship between task rule knowledge and reinforcement learning, *p* > .1 (Figure 2E).

Finally, as a test of the construct validity of reinforcement learning values, we asked whether we could replicate the phenomenon that the influence of reinforcement learning fades over time as subjects settle on rule-based response strategies^26^. Consistent with this literature, reinforcement learning was more prominent earlier in learning, *Z* = 2.0, *p* = .049, *β* = 64 (Figure 2G). In contrast, task rules influenced behavior similarly across task phases (Figure 2F), *p* > .2. Although reaction times decreased across trials, *Z* = -5.8, *p* < .001, *β* = -.084, the stability of task-rule effects on reaction times across learning suggests that the reduced influence of reinforcement learning across learning is not due to a floor effect. Moreover, the reduction in reaction times over time was accounted for when fitting our model (see *Procedures*). Learning phase effects were qualitatively unchanged when modeling trial numbers rather than learning blocks. These results demonstrate the reliable separation of rules and reinforcement learning.

### Dopamine physiology

We hypothesized that ventral striatal D2 receptors influence behavioral control. We focused our analyses on the ventral striatum because of its role in model-free and model-based learning^2^. A subset of subjects (n = 52) underwent three positron emission tomography (PET) scans to assess distinct aspects of dopamine physiology: D2/3 receptor availability with [^11^C]raclopride (raclopride has high affinity to both D2 and D3 receptors), dopamine synthesis capacity with [^18^F]fluoro-l-m-tyrosine (FMT), and dopamine release using [^11^C]raclopride displacement with methylphenidate. In the placebo condition, we found an interaction between D2/3 availability in the ventral striatum and the influence of reinforcement learning values on reaction times, *Z* = 2.3, *p* = .022, *β* = 78 (Figure 3A), with higher D2/3 availability corresponding to a reduced influence of reinforcement learning. Furthermore, higher D2/3 availability was associated with an *increased* influence of task rules on reaction times on placebo, *Z* = - 2.8, *p* = .005, *β* = -2.0 (Figure 3B). These effects are not due to fluctuations in overall task performance, as D2/3 availability was not related to average reaction times in the task nor in a separate simple reaction time task (i.e., with no rewards or predictive features), all *ps* > .1. These results show that higher D2/3 availability is associated with a shift to rule-based, rather than reinforcement-learning-based, behavioral control (Figure 3C).

**Figure 3.**
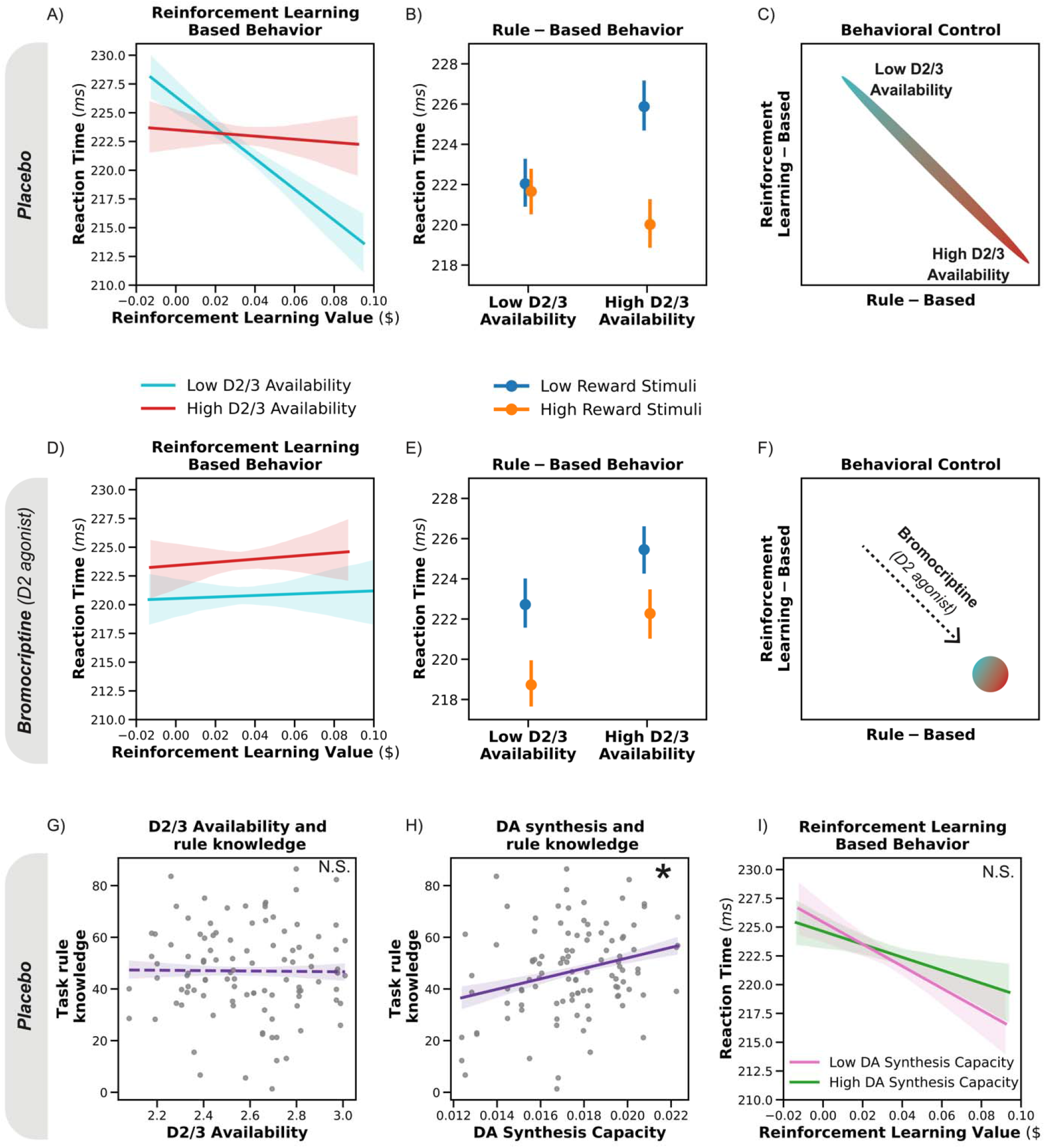
Dopamine D2/3 receptor function shifts behavioral control between reinforcement learning and rules. A-C depict placebo, and D-F depict bromocriptine. **A)** Higher D2/3 availability subjects show a reduced influence of reinforcement learning on reaction times. **B)** In contrast to *A*, higher D2/3 availability subjects show an *enhanced* influence of task rules on reaction times, as evidenced by larger reaction time differences between low and high reward stimuli. **C)** Higher D2/3 availability is associated with a shift in behavioral control from reinforcement-learning-based to rule-based. **D)** Bromocriptine, a D2 agonist, reduces the influence of reinforcement learning in subjects with lower D2/3 availability. **E)** Bromocriptine enhances the influence of task rules in subjects with lower D2/3 availability. **F)** Bromocriptine causes lower D2/3 availability subjects to favor the use of rule-based behavioral control. **G)** D2/3 availability does not relate to task rule knowledge. Task rule knowledge is operationalized as the difference in the reward probability ratings between high- and low-reward probability stimuli. **H)** Subjects with higher dopamine synthesis capacity have better task rule knowledge. **I)** Dopamine synthesis capacity does not influence reinforcement learning. *A*,*D and I depict a partial regression that removes task rule differences from reinforcement learning values. In the above panels, the median split of D2/3 availability and dopamine synthesis capacity is for visualization only, and the statistical analyses use continuous values. Error bars depict the SEM*.

To causally test whether D2 receptors influence behavioral control, we administered bromocriptine, a D2 agonist (with low D3 affinity^27^), in a placebo-controlled, double-blind study. We predicted that bromocriptine would shift behavioral control from reinforcement-learning-based to rule-based in subjects with low D2/3 availability. There was a three-way interaction between bromocriptine, D2/3 availability, and reinforcement learning value, *Z* = -2.9, *p* = .004, *β* = -140, confirming that bromocriptine blunts the influence of reinforcement learning in lower, relative to high, D2/3 availability subjects (Figure 3D). Moreover, we found a three-way interaction between bromocriptine, D2/3 availability, and task rule, *Z* = 3.1, *p* = .002, *β* = 3.0, indicating that bromocriptine enhanced rule-based responding in lower, relative to higher, D2/3 availability subjects (Figure 3E). These effects are not due to an influence of bromocriptine on overall task performance, as bromocriptine did not influence average reaction times in the task or in a separate simple reaction time task, all *ps* > .1. These results show that elevated D2 receptor activation, due to higher baseline D2/3 availability or bromocriptine administration, shifts behavioral control from reinforcement learning to task rule use (Figure 3F).

Because bromocriptine binds to D2 receptors throughout the brain, bromocriptine’s effects on behavioral control could arise due to its influence on prefrontal D2 receptors. It is also possible that our findings are not specific to D2 receptor activation, and any general stimulation of the dopamine system shifts behavioral control. As a negative control for these possibilities, we administered tolcapone to the same subjects. Unlike bromocriptine, tolcapone does not bind to dopamine receptors but inhibits dopamine clearance by disrupting COMT. Tolcapone is thought to influence dopamine clearance more strongly in the prefrontal cortex than the striatum^28–31^. If behavioral control is mediated by prefrontal dopamine tone or nonspecific dopamine system activation, tolcapone could influence task behavior similarly to bromocriptine. However, we did not find significant two-way interactions between tolcapone and either reinforcement learning or rule use, nor three-way interactions between tolcapone, D2/3 availability, and reinforcement learning or rule use, Table S2. These results indicate a selective role for striatal D2/3 receptors in behavioral control.

Rule-based behavior requires knowledge of the task rules, and dopamine system function has been linked to executive processes that could support the formation and maintenance of task rules^32–34^. An alternative explanation of our results is that subjects with higher D2/3 availability form better knowledge of task rules, resulting in a larger influence of rules on reaction time. Because we measured subjects’ explicit knowledge of the reward probabilities associated with the stimuli, we could distinguish the formation of task rule knowledge from the influence of rule knowledge on reaction times. We found no relationship between D2/3 availability and task rule knowledge (Figure 3G).

In contrast, dopamine synthesis capacity, which has been linked to executive processing^35,36^, was positively correlated with task rule knowledge, *Z* = 3.5, *p* < .001, *β* = 2.9 (Figure 3H). Similarly, dopamine synthesis capacity was associated with an increased influence of task rules on reaction times, *Z* = -2.1, *p* = .039, *β* = -1.4. In our task, behavioral control is the balance between the expression of task rules and reinforcement learning in reaction times. Unlike with D2/3 availability, we did not find a relationship between dopamine synthesis capacity and the influence of reinforcement learning value, *p* > .1 (Figure 3I). These results reveal a double dissociation between D2/3 availability and dopamine synthesis capacity, with D2/3 availability influencing reinforcement learning but not task rule knowledge and dopamine synthesis capacity showing the opposite relationship. Finally, we did not find evidence for a relationship between dopamine release, or DRD2 or COMT genotype on the influence of task rules on reaction times (see *Supplemental Results*).

### Dopaminergic influence on frontostriatal connectivity

What is the mechanism by which D2/3 receptors influence behavioral control? Striatal D2 receptors gate the influence of prefrontal inputs on striatal medium spiny neurons^14,15^. Specifically, higher D2/3 receptor activation could enhance the influence of prefrontal inputs, facilitating the expression of rule-based behavior. To test this hypothesis, we assessed functional MRI data (fMRI) acquired prior to behavioral testing in each drug session from subjects with PET imaging and adequate fMRI data (*n* = 44; Figure 1A; see also *Procedures*).

Because we found an interaction between bromocriptine and ventral striatal D2/3 availability on behavioral control, we conducted a whole-brain analysis to identify brain areas where functional connectivity with the ventral striatum showed the same interaction (Figure 4A). This analysis identified a cluster in the bilateral anterior inferior frontal sulcus and frontal pole, *p* < .05, cluster-corrected threshold (Figure 4B, C). We extracted individual subject parameter estimates to visualize this interaction (Figure 4D). This analysis revealed a qualitatively similar pattern to the behavioral results: bromocriptine enhanced frontostriatal connectivity more strongly for subjects with low D2/3 availability. These results suggest that D2/3 receptor activation enhances behavioral control by increasing the strength of frontostriatal connectivity.

**Figure 4:**
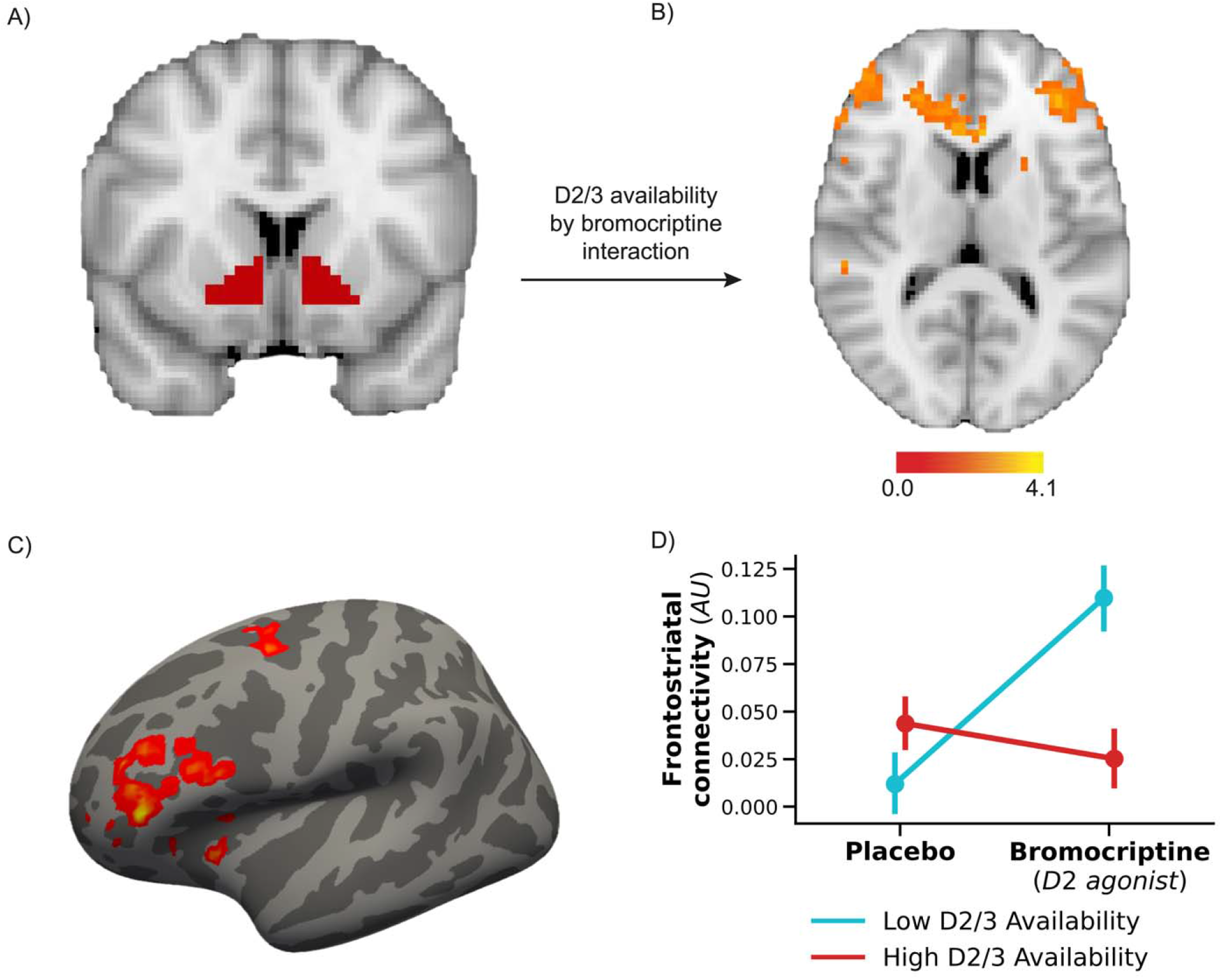
D2/3 receptors modulate ventral striatal functional connectivity with the prefrontal cortex. **A)** The ventral striatum was used as a seed region to identify brain regions where connectivity was modulated by D2/3 availability and bromocriptine, a D2 agonist. **B)** Regions within the prefrontal cortex exhibited connectivity with the ventral striatum modulated by D2/3 availability and bromocriptine. **C)** Depiction of activations in *B* on the cortical surface. Darker gray indicates sulci, and lighter gray indicates gyri. The inferior frontal sulcus and the frontopolar cortex showed connectivity modulated by D2/3 availability and bromocriptine. **D)** Visualization of the interaction in **B** and **C** from the frontopolar cortex/inferior frontal sulcus. Bromocriptine enhances connectivity between these regions and the ventral striatum more strongly for subjects with low dopamine synthesis capacity. Because this figure was derived from the group activation map, it should be interpreted as a visualization of the interaction and not an independent statistical test.

## Discussion

Our results show that activation of D2/3 receptors increases the use of rule-based over model-free knowledge without influencing the acquisition of rule knowledge. In contrast, higher dopamine synthesis capacity was linked to better explicit knowledge of the task rules and a stronger influence of that knowledge on reaction times. Importantly, dopamine synthesis capacity was unrelated to the influence of reinforcement learning. This double dissociation shows that dopamine synthesis capacity is linked to the formation and use of model-based knowledge but does not influence behavioral control.

Previous reports have found mixed evidence for a relationship between presynaptic dopamine in the striatum and model-based behavior in a two-stage decision task^37,38^. However, these studies did not separately examine the acquisition of model-based knowledge. Our results show that striatal dopamine synthesis capacity influences model-based behavior indirectly by influencing the formation of model-based knowledge rather than directly regulating behavioral control. This interpretation is consistent with recent data showing that dopamine synthesis capacity is associated with the willingness to engage in cognitive effort^19^. More broadly, our results show that dissociable components of the striatal dopamine system influence the formation of model-free and model-based knowledge and the decision about which to use.

Our neuroimaging findings showed that D2/3 receptors also influence the strength of frontostriatal connectivity, offering a mechanism by which activation of D2/3 receptors could increase behavioral control. The regulation of behavioral control by striatal D2/3 receptors may complement the role of prefrontal D1 receptors in working memory maintenance^39,40^, which is required for many model-based behaviors. It is also possible that prefrontal D2/3 receptors contribute to our results. Notably, our PET findings are specifically linked to ventral striatal D2/3 availability, and D2/3 receptors are much less strongly expressed in the prefrontal cortex than in the striatum^41,42^. Additionally, tolcapone did not influence behavioral control. Nonetheless, we cannot rule out a contribution of prefrontal D2/3 receptors, and future research is needed to disentangle the relative contributions of D2/3 receptors in the prefrontal cortex and the striatum in behavioral control.

D2 receptors may be particularly well-suited to regulating behavioral control because of their role in responding to negative feedback^9^. Our results are consistent with a model in which the D2 system responds to negative feedback by activating model-based behavioral control. The D2 system may enable habitual behavioral control when outcomes are generally positive but trigger a shift to more reliable model-based decision-making upon salient losses^4,43^. However, our study was not designed to compare the impacts of losses versus rewards, and future work is needed to test this hypothesis.

Our results show that the influence of bromocriptine on behavioral control depends on the baseline state of an individual’s D2 availability. This baseline dependence has been observed in other aspects of cognition where administration of the same drug can have opposite effects depending on individual dopamine physiology^44^. These inverted-U relationships between dopamine and cognition may help to explain inconsistent findings in previous investigations into the influence of dopamine on behavioral control. While some studies have found that administration of the dopaminergic drug L-DOPA enhances^45,46^ model-based behavior in a two-step decision-making task, another study did not find this relationship^47^. Additionally, the D2/3 antagonist amisulpride increased model-based behavior in a two-step decision-making task^6^. L-DOPA enhances dopamine release, which will increase the activation of many classes of dopamine receptors with differing influences on behavior and differing baseline expression levels. Most importantly, our results indicate an inverted-U relationship between activation of D2/3 receptors and behavioral control^36,48,49^. Therefore, modulation of D2/3 receptors could have opposing consequences for behavioral control depending on the baseline availability of D2/3 receptors.

This inverted-U relationship between D2/3 and behavioral control has relevance to clinical disorders linked to disruption in the dopamine system. Schizophrenia, which is linked to an overexpression of D2 receptors and can be treated with D2 antagonists^50^, is associated with reduced behavioral control^51^ and disrupted performance in the task used in this study^52,53^. Although the authors are not aware of data from Parkinson’s disease patients performing this task, administration of bromocriptine or other dopamine agonists in Parkinson’s disease patients can cause deficits in other forms of behavioral control, including compulsive gambling, binge eating, or overspending^54,55^.

The direction of our findings contrasts with these clinical patterns of poorer behavioral control linked to overexpression or overactivation of D2 receptors. However, an inverted-U model can account for this discrepancy^36,48,49^. Low D2/3 subjects in our study benefit from a moderate dose of bromocriptine, whereas very high D2/3 activation due to clinical state or pharmacological intervention can impair behavioral control. Ultimately, a mechanistic understanding of the neural basis of behavioral control could lead to improved treatment options for psychiatric and neurological disorders linked to dopamine system dysfunction.

## Methods

### Procedure Overview

Drug and genotype procedures have been previously described^56^, and key details are reproduced here. Individuals meeting the advertised inclusion criteria were invited to the Helen Wills Neuroscience Institute at the University of California, Berkeley, to provide a saliva sample for genotyping. Subjects meeting medical and genotype criteria (see below) were scheduled for three pharmacological study sessions to be completed on different days. Between 2 to 86 days separated subsequent sessions (median = 7, mean = 10.2, SD = 9.7). At each of the three sessions, subjects were administered a single dose of bromocriptine, tolcapone, or placebo, after which they performed a cognitive task in an MRI scanner, the results of which have been previously reported ^56^.

Following the scanner task and outside of the scanner, they performed the probabilistic response time task described here. Although overall session start times varied between 7 a.m. and 4 p.m., efforts were made to keep start times consistent across sessions for each subject. A subset of the subjects performed two PET sessions. All subjects gave written, informed consent in accordance with the procedures approved by the Committee for the Protection of Human Subjects at the University of California, San Francisco, the University of California, Berkeley and the Lawrence Berkeley National Laboratory and were compensated for their participation.

#### Subjects

Healthy young subjects were recruited for genetic sampling from the University of California, Berkeley, community, and surrounding area using online and print advertisements. Subjects affirmed at the time of genetic sampling that they met initial inclusion criteria: (1) 18–30 years old, (2) right-handed, (3) current weight greater than 100 pounds, (4) able to read and speak English fluently, (5) nondrinker or light drinker (women: <7 alcoholic drinks/week; men: <8 alcoholic drinks/week), (8) no recent history of substance abuse, (9) no history of neurological or psychiatric disorder, (10) not currently using psychoactive medication or street drugs, (11) not pregnant, and (12) no contraindications to MRI (e.g., no claustrophobia, pacemakers, history of seizures, or MRI-incompatible metal in body).

Genetically eligible subjects underwent a medical screening with an on-site physician or nurse practitioner and a liver function test to ensure there were no medical contraindications to tolcapone and bromocriptine use and to verify the absence of neurological and psychiatric history. Seventy-seven subjects completed the behavioral study, not including three who dropped out. We did not exclude any subjects from the behavioral analyses. One subject did not perform the bromocriptine session, but their placebo and tolcapone data are included.

### Genotype

Saliva samples were obtained using Oragene collection kits with stabilizing liquid (DNA Genotek). Genotyping of COMT (rs4680) and Taq1A (rs1800497) SNP testing was performed at the UCSF Genomics Core, Vincent J. Coates Genomics Sequencing Laboratory, and Kashi Clinical Laboratories using polymerase chain reaction-based TaqMan technology (Applied Biosystems). Only individuals who were homozygous for either the Val or Met allele of the COMT polymorphism were invited to participate in the remainder of the study. Taq1A genotypes were binned according to the presence (“A1+”) or absence (“A1−”) of any copies of the A1+ minor allele. Subjects were selected based on compound COMT/Taq1A genotype, with roughly equal representation in each of the following groupings: Met/A1+ (n = 20), Val/A1+ (n = 17), Met/A1− (n = 21), Val/A1− (n = 19).

#### Drugs

During each session, subjects received a single oral dose of bromocriptine (1.25 mg), tolcapone (200 mg), or placebo. These doses were selected based on their established efficacy in eliciting changes in cognitive performance^33,57–59^. After administration, bromocriptine reaches peak plasma concentrations between 0.5 and 3.5 hr (mean time to peak = 1.7 hr) and has an elimination half-life of 3–7 hr^60,61^. Tolcapone reaches peak plasma concentration, on average, 1.8 hr after oral administration and has an elimination half-life of about 2 hr^62^. Subjects began the task approximately 3.1 hr after drug/placebo administration, after the most probable peak concentration time but within the elimination half-life of both drugs.

The order of drug administration was double-blind and counterbalanced across subjects (and within genotype groups). After each session, subjects were asked whether they received a placebo or a drug that day. As a group, subjects demonstrated no better than chance-level accuracy in their guesses, and rates of “drug” vs. “placebo” guesses did not differ significantly across the three sessions, χ2(2) = 3.2, *p* = .2 (excludes four omitted responses).

#### PET procedures

Fifty-two of the subjects completed three PET scans. The PET procedures and analysis techniques are described in prior reports using this dataset^63^. Briefly, subjects underwent an [^18^F]fluoro-l-m-tyrosine (FMT) PET scan to measure dopamine synthesis capacity, a [^11^C]raclopride PET scan to measure D2/3 receptor occupancy, and a [^11^C]raclopride PET scan one hour following a 30mg oral dose of methylphenidate to measure dopamine release. All PET data were acquired using a Siemens Biograph Truepoint 6 PET/CT scanner. Data were reconstructed using an ordered subset expectation maximization algorithm with weighted attenuation, corrected for scatter, smoothed with a 4 mm full width at half maximum (FWHM) kernel, and motion corrected. Ventral striatal ROIs were hand-drawn on subjects’ T1-weighted images according to established procedures^64^. FMT data were analyzed using a reference region Patlak model^65^ to determine net tracer influx (K_i_) as the outcome variable of interest reflecting dopamine synthesis capacity. Raclopride analyses used a reference tissue reversible tracer mode^66^ to calculate non-displaceable binding potential (BP_ND_) as the outcome of interest reflecting receptor availability.

#### MRI procedures

Functional and anatomical MRI data was obtained during each drug session with a Siemens 3T Trio Tim scanner at UC Berkeley’s Brain Imaging Center. Of the fifty-two subjects with PET data, one subject did not complete the placebo MRI session, three subjects had unusable MRI acquisitions, and four subjects were excluded for poor scanner task performance. This resulted in a sample of forty-four subjects with PET, fMRI, and behavioral data. After preprocessing and regressing task and noise effects from the task-fMRI data^67,68^ (*Supplemental Methods*), we performed functional connectivity analyses using a multilevel model in FSL 6.0.3. At the individual run level, we created a model using the average signal from the same ventral striatal mask used in the PET analyses, the whole-brain signal, and a regressor containing censored volumes^69^. At the subject level, we modeled the effects of both drugs on ventral striatal functional connectivity. At the group level, we modeled the effects of ventral striatal D2/3 availability and its interaction with the drug condition on ventral striatal functional connectivity. Z-statistic images were thresholded non-parametrically using clusters determined by Z>1.7 and a (corrected) cluster significance threshold of P=0.05^70^. To visualize the activation clusters on the surface mesh, we used RF-ANTs to register results to *fsaverage* surface coordinates^71^.

#### Task design

Subjects completed a speeded reaction time task with probabilistic rewards known as the *salience attribution task*^*52*,*53*^. Each trial began with a stimulus presented on both the left and right sides of the screen. After a variable interval, a central target appeared. Subjects were instructed to respond to the target by pressing a key as quickly as possible.

Subjects were not told whether a trial was rewarded or unrewarded. However, the pre-target cues provided predictive information. There were four cue stimuli that could have one of two colors and one of two shapes. One feature predicted a reward with an 85% probability (e.g., “pink” in Figure 1B), and the other feature of that dimension (e.g., “green”) predicted a reward with a 15% probability. The other dimension (e.g., “shape”) did not predict reward. To facilitate subjects’ use of rules, these probabilities were fixed for the task’s 128 trial duration. The reward-predictive dimension was balanced across subjects and drug conditions. To prevent interference across sessions, new shapes and colors were used in each drug session and counterbalanced across subjects. We refer to the two stimuli with the reward predictive feature as “high-reward stimuli” and the other two stimuli as “low-reward stimuli.”

On rewarded trials, subjects earned more monetary reward for faster responses, provided they responded faster than a predetermined threshold. This threshold was set individually and for each session based on a simple RT task immediately preceding the salience attribution task. In the simple RT task, subjects responded to the target for 35 trials without any informative cues and received no feedback or rewards. Monetary rewards were accompanied by a coin sound. If responses on reward trials were too slow, or subjects responded before the target appeared, they were given feedback (“Miss” or “Too Early,” accompanied by a “beep” sound). However, they earned a small ($.02) reward to ensure equal reward probabilities across subjects and sessions. On unrewarded trials, subjects received no feedback about their performance and no monetary reward.

Subjects were also asked to explicitly report the reward probability of each of the four stimuli using a slider that ranged from 0 to 100%. Subjects made these judgments once halfway through the task and again at the end of the task.

#### Data analysis

Behavioral data were analyzed using mixed linear models implemented in statsmodels 0.11.0. Drug conditions were treatment-coded with the placebo condition as the baseline. All other effects were deviation coded. PET values were z-scored prior to inclusion in models. Although some figures show PET data as median split for visualization purposes, all statistical analyses were conducted on continuous data. All

RL analyses included reinforcement learning value and task rule as regressors, as well as their two and three-way interactions with drug and PET measures in the same model. All mixed-effects models contained random intercepts for each subject. We chose this random effects approach because theoretical and modeling work shows that mixed-effects models generalize most effectively when they use the maximal random effects structure that is justified by the design and does not create convergence failures^72^. Random slopes caused convergence failures and were removed from our models. Analysis of the simple RT task preceding the main task included trial number as a nuisance covariate.

Due to a computer error, reaction times slower than the threshold time were not recorded, which occurred in 9.6% of trials. Nevertheless, these trials contain information because we know that subjects responded more slowly than the response threshold. Therefore, we implemented a data imputation procedure to estimate these reaction times (Supplemental Methods). We note that this procedure did not change the qualitative patterns in the reported results.

#### Reinforcement learning model

We used reaction time data to infer learning in the task, an approach used successfully in similar tasks^73–75^. The key concept of the model is that trials with higher values should be associated with faster reaction times. We modeled reaction times as a function of reinforcement learning value using linear regression. We quantitatively compared several model variations (*Supplemental Methods*), and the best-fitting model is described here. The value of each stimulus was defined as the sum of the weights of the features comprising that stimulus:

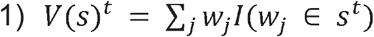

Where *V(s)*^*t*^ is the value of stimulus *s* on trial *t*, the *w*_*j*_ are the weights of the different features (e.g., square, circle, green, purple), and 1 is an indicator function equal to one if the feature was present on that trial and 0 otherwise. This model learns the values of the individual features and computes the stimulus value as the sum of the component feature values. The model generalizes information learned about individual features (e.g., “square”) across different stimuli (“purple square” and “green square”), accurately reflecting the underlying reward structure of the task.

After feedback, the weights of features present on that trial are updated according to the Rescorla-Wagner rule:

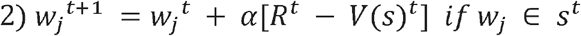

Where R^t^ is the reward earned on trial *t* and α, the learning rate, is determined separately for rewards and reward omissions^76^:

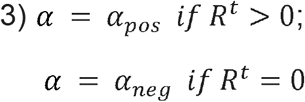

Finally, to reflect memory decay^77^, weights of features not present on a trial are decremented by a free parameter *d*,

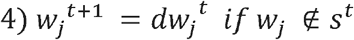

We jointly fit the parameters of the regression model and the parameters of a reinforcement learning model using Scipy’s minimize function. We calculated likelihoods from the regression fits using:

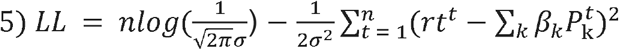

Where *n* is the number of trials, the 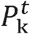 are different predictors of RT at trial *t*, and *β*_*k*_ are regression coefficients. In addition to value, we included three other mean-centered predictors of RT (*P*_*k*_) in the model: 1) an indicator function of whether the last trial was rewarded (*P*_*rew-last-trial*_; to account for post-reward slowing)^78,79^, 2) a linear function of trial number within a block (*P*_*trial*_), and 3) a main effect of the learning block (*P*_*block*_). The latter two regressors isolate value-related changes in RT from stimulus-general improvement in RTs over time.

We fit the data of all subjects and sessions simultaneously to compute parameters of the reinforcement learning model (*α*_*pos*_, *α*_*neg*_,*d, β*_*value*_,*β*_*rew-last-trial*_,*β*_*trial*_,*β*_*block*_; Table S1) and used these parameters to generate reinforcement learning value estimates for every trial. Pooling subjects has a regularizing effect on parameter estimates that helps to avoid over-fitting and improve parameter identifiability (See *Parameter recovery study* in *Supplemental Methods)*^*2*,*24*,*74*^. Pooling over sessions ensures that differences observed between drug conditions are not due to biased model-fitting to an individual condition. Thus, our modeling approach was designed to deliver robust estimates of trial-by-trial value, enabling us to assess how dopaminergic drugs and PET measures influence behavioral control.

To assess whether reinforcement learning and task rules contribute to behavior in the task (Figure 2C), we used leave-one-subject-out cross-validated model-fitting. For each subject, we fit the parameters of the reinforcement learning model to group data composed of all the other subjects. We used these parameters to generate predicted reinforcement learning values for that subject. This approach enables a statistically independent test of whether reinforcement learning values influence reaction times. Having established this effect, we used the reinforcement learning values described above (which use the same set of parameters for all subjects) for subsequent analyses.

## Data Availability Statement

Analysis code is available at https://github.com/iancballard/dopamine_behavioral_control.

