## Supplemental Text for "A dopaminergic basis of behavioral control"

**Supplemental Methods**

*Data Imputation*

Due to a computer error, response times slower than the response time threshold were not recorded in 9.6% of trials. This did not impact subjects’ experience of the task. Because the response time threshold was variable across trials, miss trials tended to occur when the response threshold was very fast (median threshold on miss trials was 244 ms compared to 269 ms on other trials). On trials where the response threshold was faster than the average RT, we imputed the subject’s average RT from the session. Otherwise, on trials where the threshold was slower than the average RT, we imputed the response threshold. This procedure incorporates the information that subjects were at least as slow as the threshold on missed trials when the threshold was slower than their average RT.

*Reinforcement Learning Model Comparison*

Our model comparison aimed to identify a model that best explains trial-wise variations in stimulus value that arise due to reinforcement learning. To do so, we fit and quantitatively compared four elaborations of a basic model, which we refer to as the *Single Learning Rate* model*.* Because we found a transition from reinforcement learning to task-rule response patterns over learning, we created a *Collapsing Learning Rate* model (free parameters {$\alpha$,*c*}) in which the learning rate exponentially declines over trials.

1) $\alpha^{t} = {\alpha e}^{-ct}$

Where $\alpha^{t}$ is the learning rate at trial *t*, and *c* is a free parameter that controls the steepness of the decline. This model was meant to capture a gradual transition in which rewards exert a smaller and smaller influence on value estimates as time progresses. However, this model provided a poorer fit to the data than the *Single Learning Rate* model and was not considered further.

We next constructed the *Decay* model (free parameters {$\alpha$,*d*}), which decrements the value of features that have not been recently encountered, reflecting memory decay^1^. In this model, weights of features not present on a trial are decremented by a free parameter *d*:

2) ${w_{j}}^{t+1} ={{dw}_{j}}^{t} if w_{j} \notin s^{t}$,

where *d* is bound on [0,1].

We also considered the possibility that participants were differentially sensitive to rewards and reward omissions^2^. We fit a *Dual Learning Rates* model with separate learning rates, $\alpha$_pos_ and $\alpha$_neg_, for rewards and reward omissions. We replaced equation 2 with:

3) $\alpha= \alpha_{pos} if R^{t}>0$;

$$\alpha= \alpha_{neg} if R^{t}=0$$

Finally, we fit a *Dual Learning Rates with Decay* model (free parameters {$\alpha$_pos_,$\alpha$_neg_,*d}*) with both factors in a single model. We compared models by approximating the model evidence of each model relative to the *Single Learning Rate Model* using the Aikake Information Criterion (*AIC)* to balance model-fit and model complexity. We found that the *Dual Learning Rates with Decay* provided the best account of the reaction time data (Figure S1) and used this model in subsequent analyses.

*Parameter Recover Study*

We conducted a parameter recovery study to assess the identifiability of the parameters in our model^3^. We used the best-fit parameters (Table S1) to invert our model and simulate reaction times. Second, we added normally distributed noise to these simulated reaction times with variance equal to the noise observed in the empirical data. Finally, we fit our model to the simulated reaction times. We ran 500 simulations, and the median estimated parameters are reported in Table S1. The recovered parameters were very close to the ground truth parameters, indicating that parameter estimates from our model are robust for the parameter values found in our study.

*MRI procedures*

Anatomical images were obtained using a T1-weighted magnetization prepared rapid gradient-echo (MPRAGE) sequence repetition time (TR) = 2300 ms; echo time (TE) = 2.98 ms; flip angle (FA) = 9°; bandwidth = 238 Hz/Pixel; matrix = 240 × 256; field-of-view (FOV) = 256 cm; sagittal plane; voxel size = 1mm^3. Functional MRI data was obtained while subjects performed a cognitive task described elsewhere[^4^. fMRI data was obtained over three 11-minute runs with a gradient-echo echo-planar imaging (EPI) sequence (TR = 2,000ms, TE = 24ms, flip angle = 65 degrees, field of view 224mm, 36 slices, AC-PC, voxel size = 3.0x3.0x3.5mm), 336 volumes. Functional and anatomical imaging data was processed using fMRIPrep^5–7^. Anatomical data was N4 bias-field corrected using N4BiasFieldCorrection^8^ and normalized to MNI space using nonlinear registration with antsRegistration (ANTs 2.2.0). BOLD data was motion-corrected and co-registered to the T1w reference using flirt (FSL 5.0.9)^9^ with boundary-based registration^10^. BOLD data were resampled to MNI space by concatenating the BOLD-to-T1w and the T1w-to-MNI transformations.

For the functional connectivity analysis, we created “pseudo-resting state data” by residualizing task regressors from data obtained during the cognitive task. This approach has been shown to result in similar measures of functional connectivity as traditional resting state data^11,12^. Task-related regressions were constructed for each event using a spline basis set model. Distractor, switch, ambiguous, first, error, and miss onsets were modeled using CSPLINE in AFNI^13^. Motion traces, motion derivatives, and 0th-4th order trends were included in the model. Residuals were calculated using 3dDeconvolve in AFNI. We did not apply spatial smoothing to the data.

**Supplemental Results**

We performed exploratory analyses to investigate whether our genetic measures relate to behavioral control. We did not find relationships between DRD2 or COMT alleles on the influence of task rules or reinforcement learning on behavior, *p* > .1 (Table S4 and S5).

We did not find evidence that dopamine release, as measured with [^11^C]raclopride displacement with methylphenidate, related to the influence of task rules on reaction times. However, dopamine release was related to the influence of reinforcement learning values on reaction times, *Z* = 2.1, *p* = .035, *β* = 74, but this effect was not modulated by bromocriptine, p > .1 (Table S6).

**Supplemental Figures and Tables**

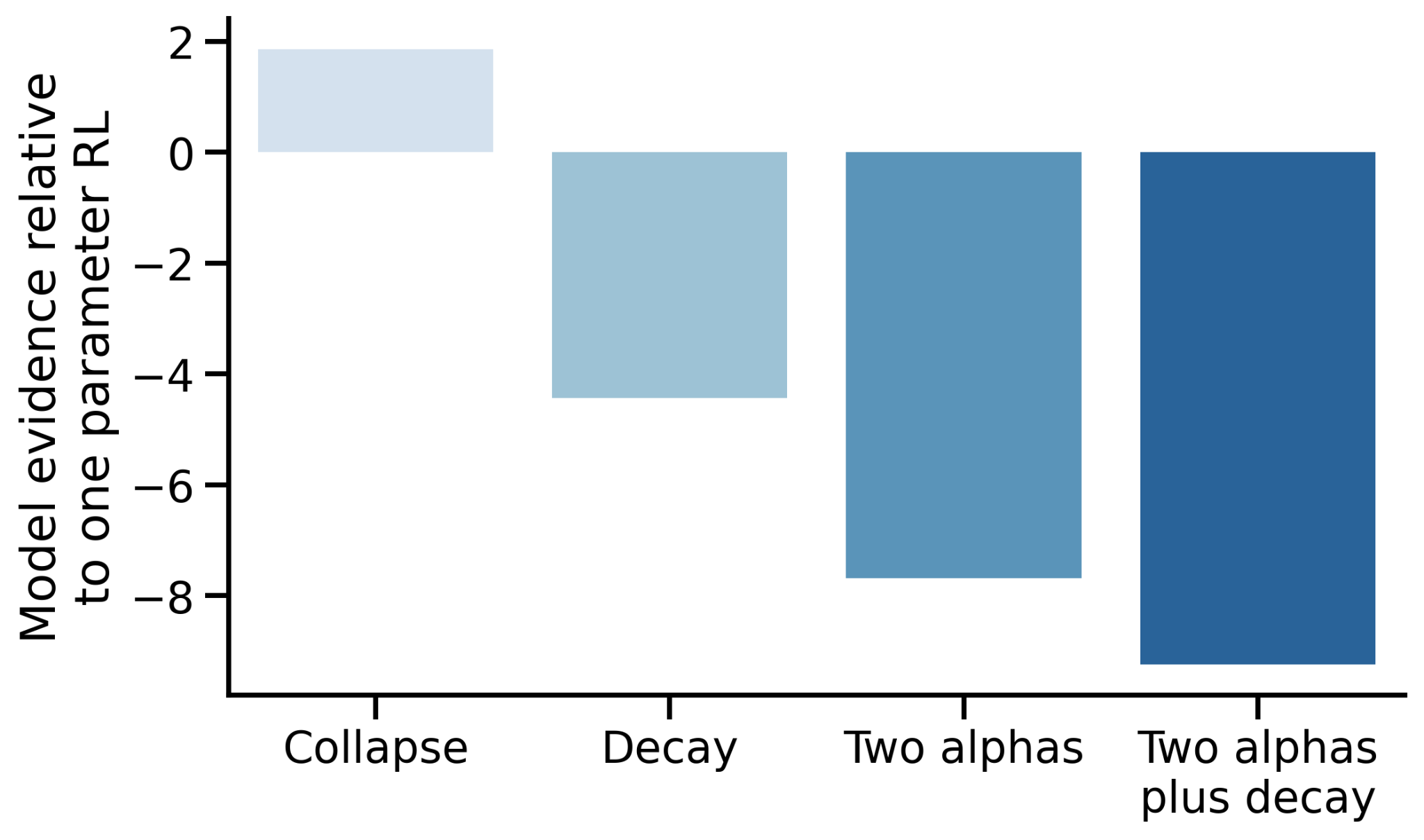

**Figure S1**: *Model comparison results*. The y-axis depicts AIC relative to one-parameter reinforcement learning. The *Collapse* model, which has a learning rate that declines exponentially over trials, performed worse than the baseline model. The *Decay* model, in which values of unseen features decline over trials, and the *Two Alphas* model, in which separate learning rates were fit for rewards and reward omissions, outperformed the baseline model. The winning model incorporated both a decay term and two learning rates.

|  | **Reward learning rate** ($\alpha_{pos}$) | **Reward omission learning rate** ${(\alpha}_{neg}$) | **Decay** (*d*) | **RL Beta** ($\beta_{value}$) | **Reward Last Trial Beta** ${(\beta}_{rew-last-trial})$ | **Trial Number Beta** ($\beta_{trial}$) | **Block Number Beta** ($\beta_{block}$) |
| --- | --- | --- | --- | --- | --- | --- | --- |
| Maximum likelihood estimate | 0.13 | 0.04 | 0.97 | -2.05 | 1.11 | -127 | -0.55 |
| Recovery study median | 0.12 | 0.04 | 0.98 | -2.03 | 1.11 | -127 | -0.58 |
| Recovery study 95% CI | [0.05,  0.19] | [0.01,  0.11] | [0.092,  1.0] | [-2.45,  -1.63] | [0.53,  1.56] | [-229,  -91] | [-1.06,  -0.06] |

**Table S1**: Maximum likelihood estimate parameters for the winning model and parameter recovery study results.

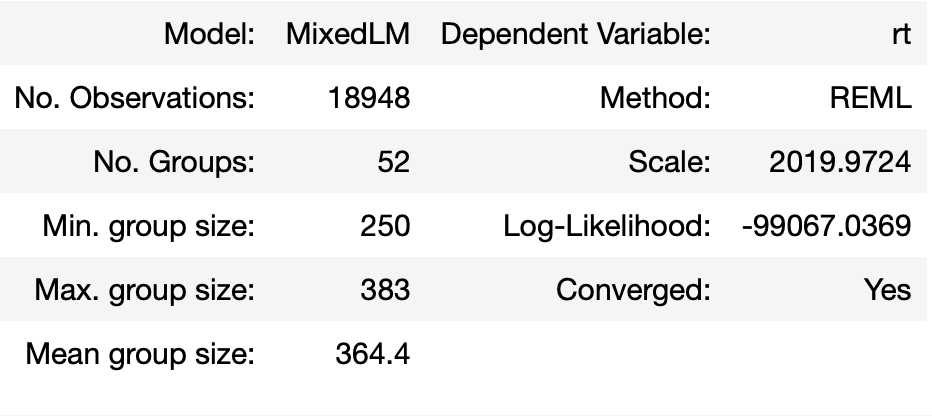

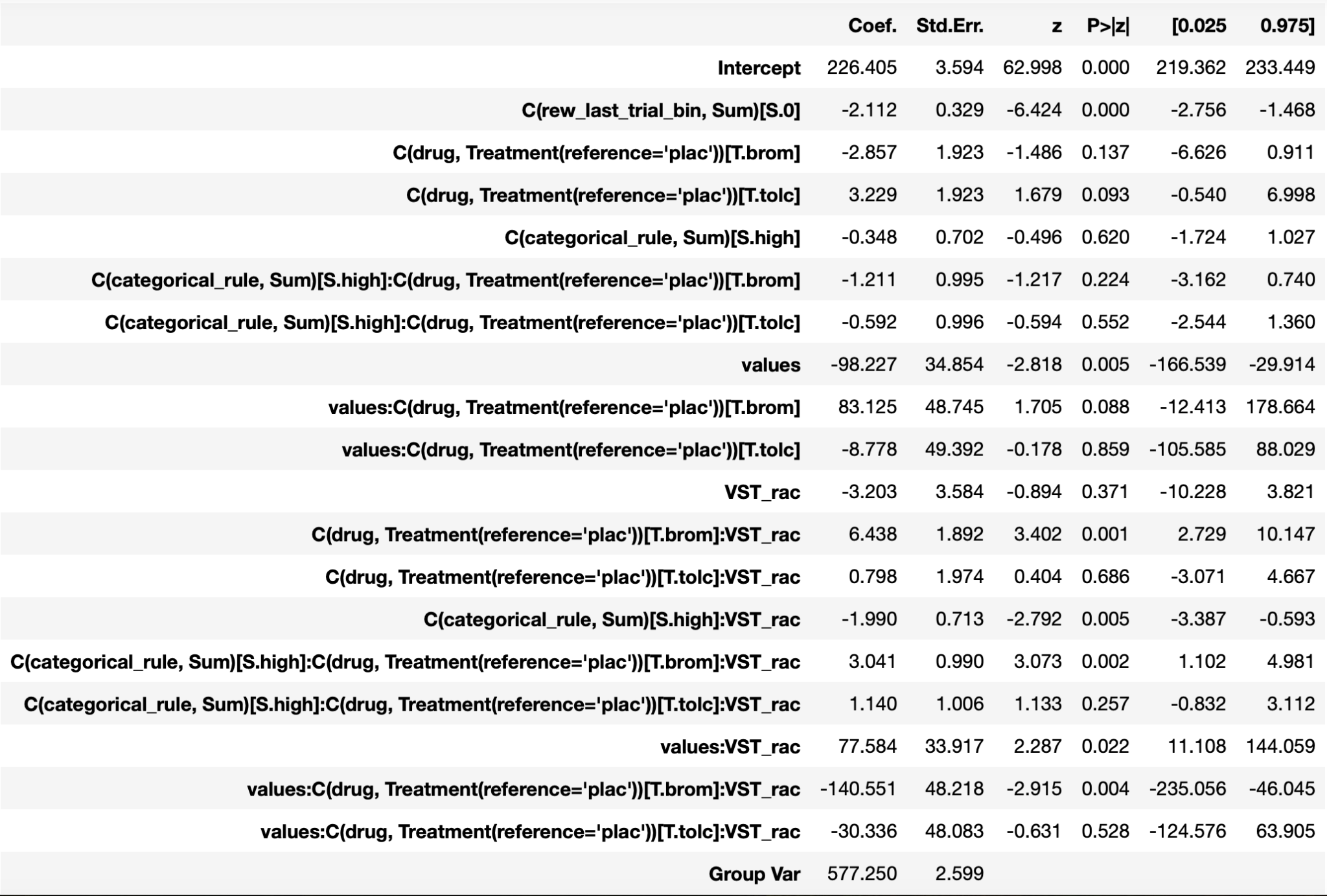

**Table S2:** *D2/3 availability and drug influence on reinforcement learning and categorical rule use.* Reaction times are the dependent variable *‘plac’ = placebo, ‘brom’ = bromocriptine, ‘tolc’ = tolcapone, ‘values’ = reinforcement learning values,* ‘VST_rac’ = ventral striatal [11C]raclopride, *‘Sum’* *refers to deviation coding, ‘T’ refers to treatment coding*.

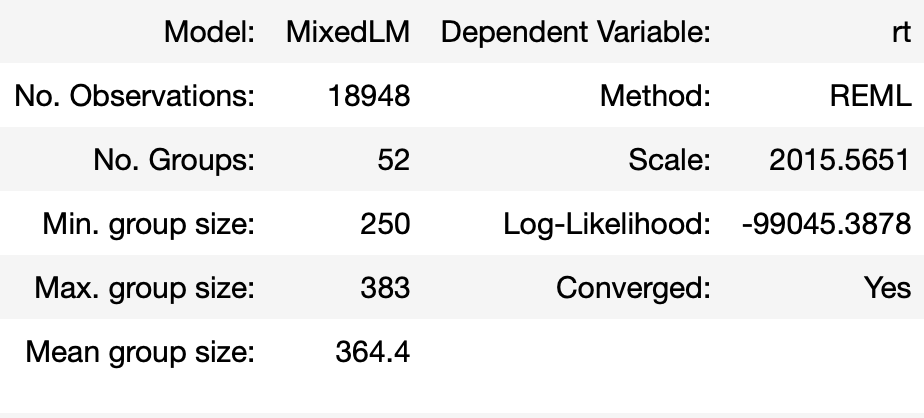

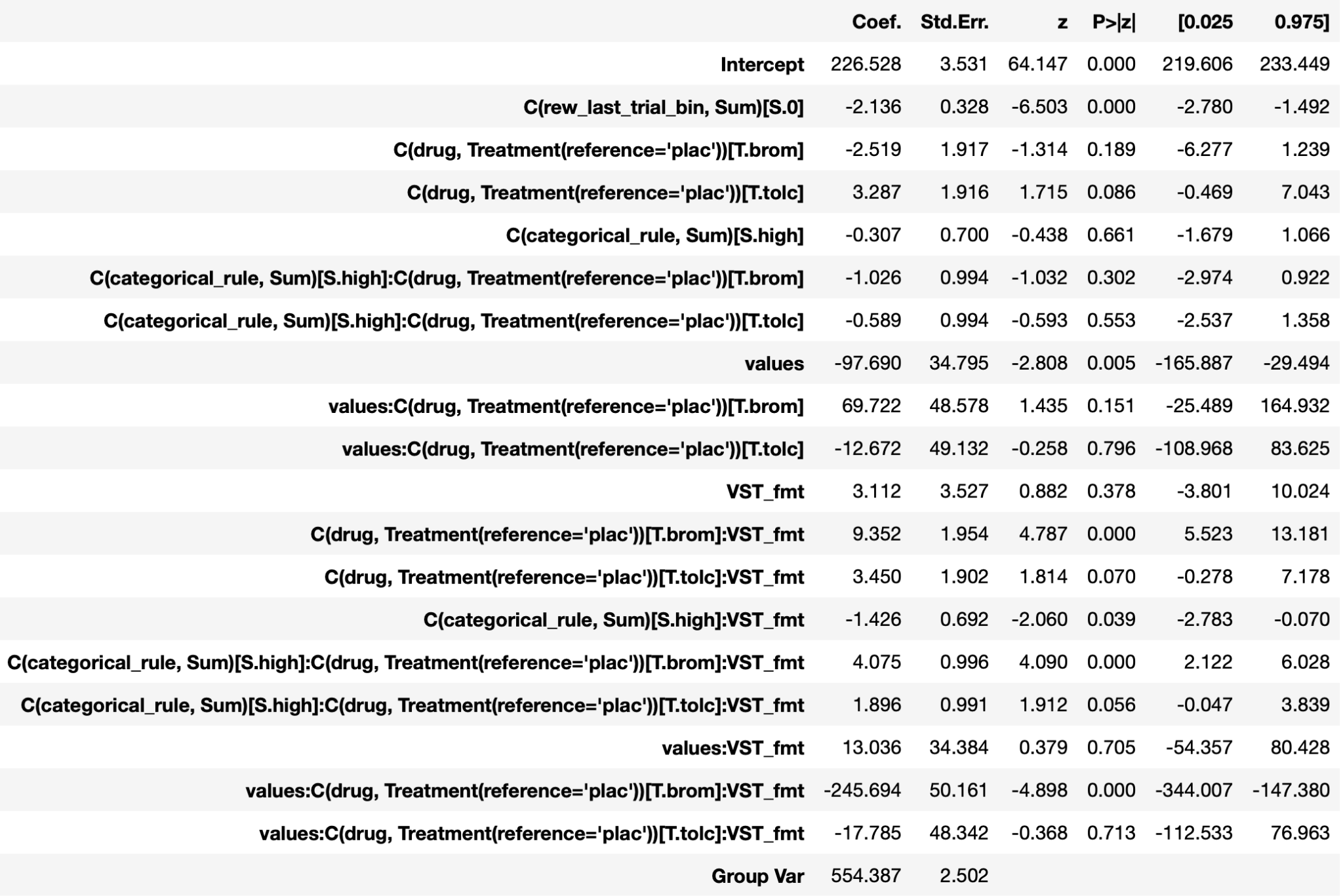

**Table S3:** *Dopamine synthesis capacity and drug influence on reinforcement learning and categorical rule use*. Reaction times are the dependent variable *‘plac’ = placebo, ‘brom’ = bromocriptine, ‘tolc’ = tolcapone, ‘values’ = reinforcement learning values,* ‘VST_fmt’ = ventral striatal [18F]FMT, *‘Sum’* *refers to deviation coding, ‘T’ refers to treatment coding*.

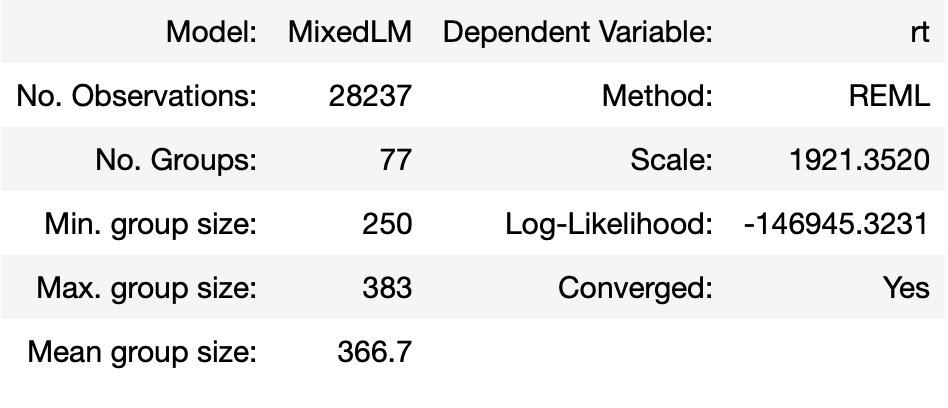

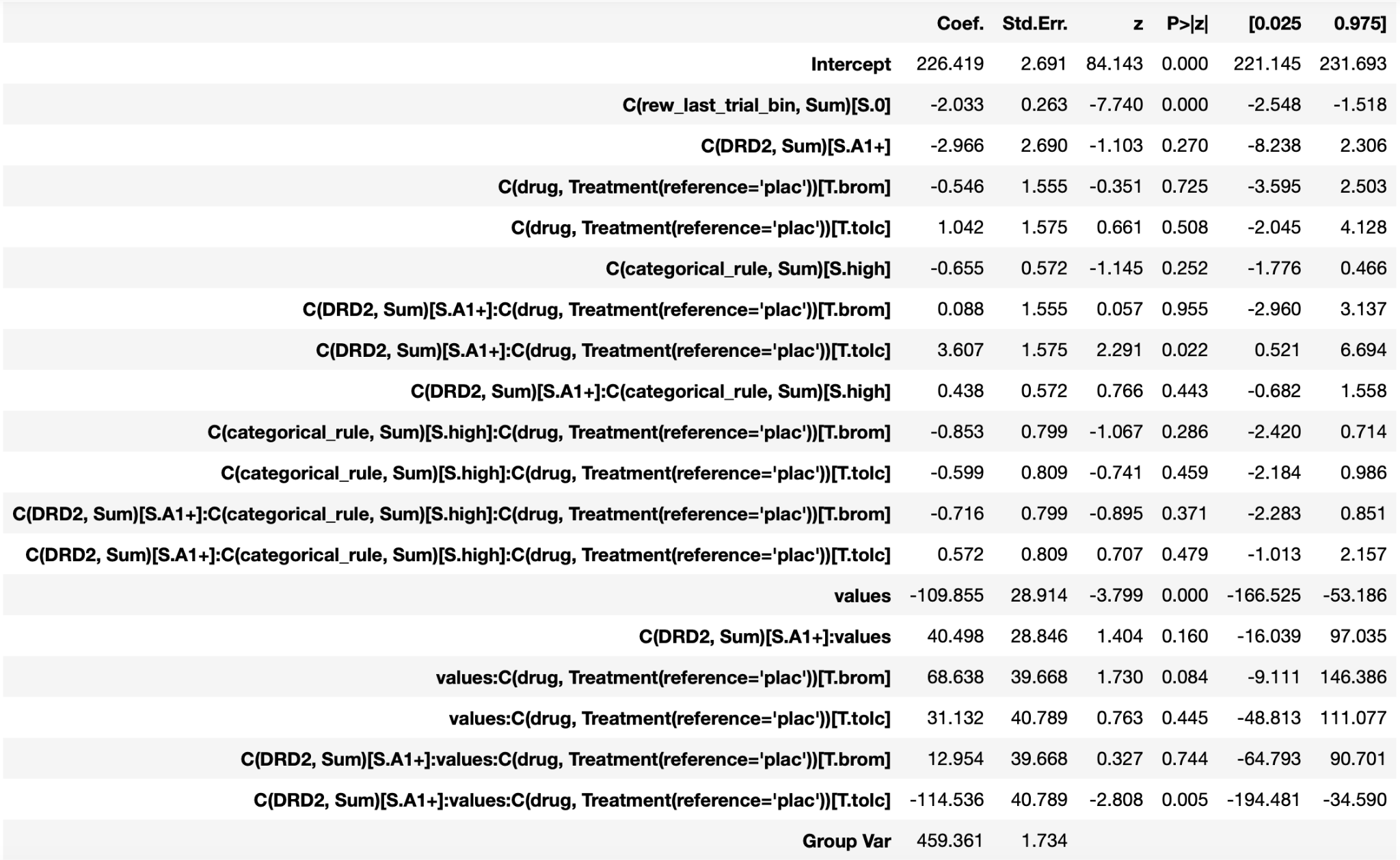

**Table S4:** *DRD2 influence on reinforcement learning and categorical rule use*. Reaction times are the dependent variable. *‘plac’ = placebo, ‘brom’ = bromocriptine, ‘tolc’ = tolcapone*, *‘values’ = reinforcement learning values, ‘Sum’* *refers to deviation coding, ‘T’ refers to treatment coding*.

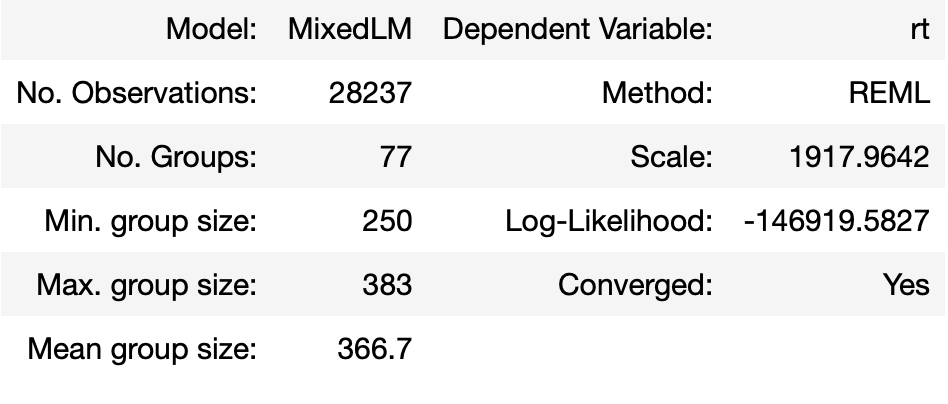

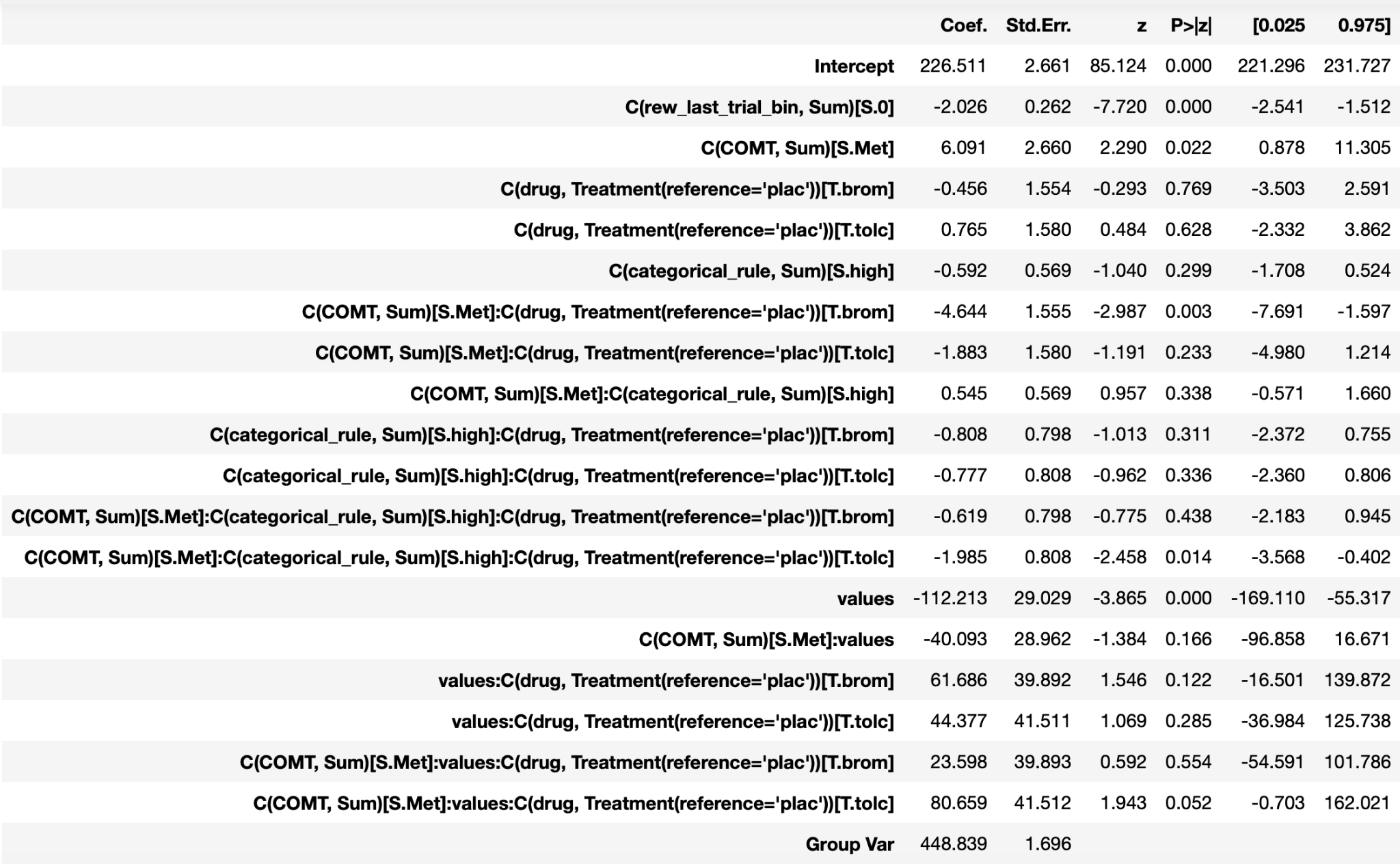

**Table S5:** *COMT influence on reinforcement learning and categorical rule use*. Reaction times are the dependent variable *‘plac’ = placebo, ‘brom’ = bromocriptine, ‘tolc’ = tolcapone, ‘values’ = reinforcement learning values, ‘Sum’* *refers to deviation coding, ‘T’ refers to treatment coding*.

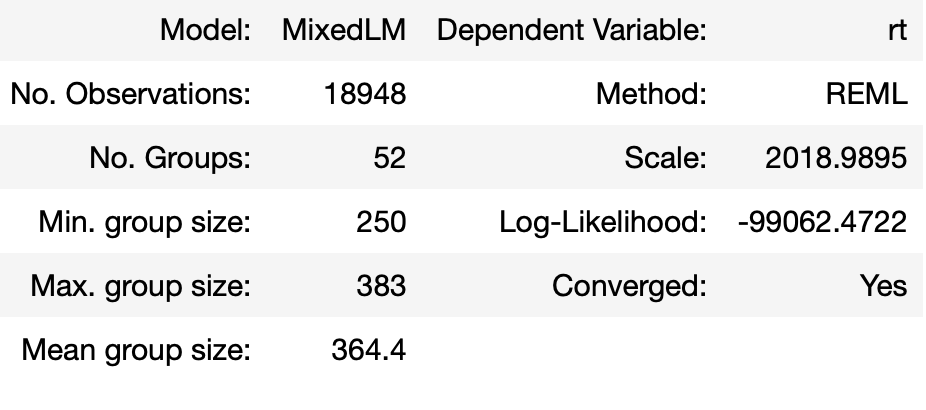

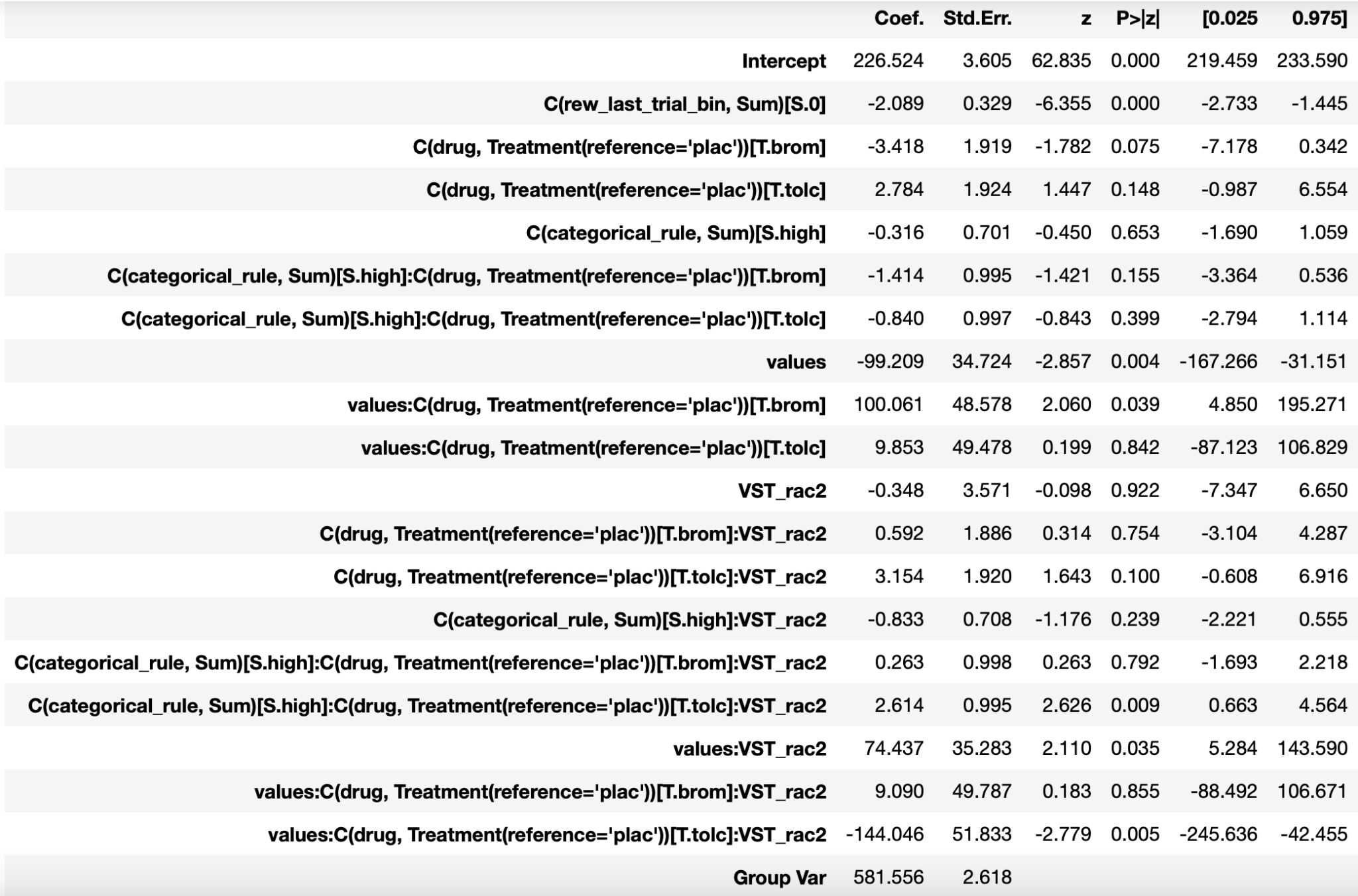

**Table S6:** *Dopamine release and drug influence on reinforcement learning and categorical rule use*. Reaction times are the dependent variable *‘plac’ = placebo, ‘brom’ = bromocriptine, ‘tolc’ = tolcapone, ‘values’ = reinforcement learning values,* ‘VST_rac2’ = dopamine availability derived from the methylphenidate challenge, *‘Sum’* *refers to deviation coding, ‘T’ refers to treatment coding*.
